# Clonal transmission and new mechanism of resistance to trimethoprim-sulfamethoxazole in *Stenotrophomonas maltophilia* strains isolated in a neonatology unit at Antananarivo, Madagascar, deciphered by whole genome sequence analysis

**DOI:** 10.1101/696765

**Authors:** Mamitina Alain Noah Rabenandrasana, Volasoa Andrianoelina, Melanie Bonneault, Perlinot Herindrainy, Benoit Garin, Sebastien Breurec, Elisabeth Delarocque-Astagneau, Zafitsara Zo Andrianirina, Vincent Enouf, Bich-Tram Huynh, Lulla Opatowski, Jean-Marc Collard

**Affiliations:** Experimental Bacteriology Unit, Institut Pasteur, Antananarivo, Madagascar; Biostatistique, Biomathématique, Pharmaco-épidémiologie et Maladies Infectieuses (B2PHI), Institut Pasteur, Inserm, Université de Versailles–Saint-Quentin-en-Yvelines (UVSQ), Paris, France; Epidemiology and Clinical Research Unit, Institut Pasteur, Antananarivo, Madagascar; Laboratoire Immuno-Hématologie CHU, CHU Pointe-à-Pitre, Abymes, Guadeloupe, France; Faculté de médecine, Institut Pasteur de la Guadeloupe, Pointe-à-Pitre, Guadeloupe; Pediatric Ward, Centre Hospitalier de Soavinandriana, Antananarivo, 97159, Madagascar; Pasteur International Bioresources network (PIBnet), Plateforme de Microbiologie Mutualisée (P2M), Institut Pasteur, Paris, France

## Abstract

*Stenotrophomonas maltophilia* has been recognized as an emerging multidrug resistant organism in hospital settings due to its resistance to a broad range of antimicrobial agents. These include β-lactams and aminoglycosides, afforded by the existence of intrinsic and acquired resistance mechanisms. Trimethoprim/sulfamethoxazole (SXT) is recommended as one of the best treatment choices against *S. maltophilia* infections; however increasing resistance to SXT has complicated the treatment. From July 2014 to March 2015, individuals and surfaces from a neonatology ward in Antananarivo, Madagascar, were longitudinally followed to assess the transmission of bacteria resistant to antibiotics between neonates, individuals (parents and nurses) and ward environments. Four *S. maltophilia* strains were successively isolated from a water-tap (N=1), from feces obtained from a newborn (N=1), and nursing staff (N=2). Antimicrobial susceptibility testing and whole genome sequencing were performed on each isolate. Based on coregenome alignment, all strains were identical and belonged to the new sequence type ST-288. They were resistant to trimethoprim-sulfamethoxazole, carbapenems and intermediate to levofloxacin. Each isolate carried the *aadB, strA, strB* and *sul1* genes located in a class I integron but variants of the *dfrA* gene were absent. We assessed by PROVEAN analysis the single nucleotide mutations found in *folA, folC* and *folM* genes and only the mutation in *folA* (A114T:GCC→ACC) has an effect on the activity of trimethoprim. Our findings demonstrated the prolonged presence of SXT-resistant *S. maltophilia* in a clinical setting with consecutive transfers from the environment to a newborn and staff based on the isolation dates. We also hypothesized that single nucleotide mutations in *folA* could be responsible for trimethoprim resistance.

## INTRODUCTION

*Stenotrophomonas maltophilia* is a non-fermentative Gram-negative bacterium generally found throughout the environment (soil, sewage, plants). It is also occasionally isolated in hospitals where this bacterium is currently regarded as an important opportunistic pathogen. It causes a large range of clinical syndromes such as bacteraemia, sepsis, pneumonia, meningitis, endocarditis, septic arthritis, urinary infections, and endophthalmitis (1, 2), especially in hospitalized immunocompromized patients or patients with underlying disease. *S. maltophilia* has been recognized as one of the leading nosocomial multidrug resistant organisms due to its resistance to a broad range of antimicrobial agents, including β-lactams and aminoglycosides, afforded by the existence of intrinsic and acquired resistance mechanisms (3). However it remained susceptible to fluoroquinolones, polymyxins and trimethoprim/sulfamethoxazole (SXT) (4).

SXT in association with ticarcillin and clavulanic acid is traditionally recommended as one of the first choices against *S. maltophilia* infections but fluoroquinolones (also in association with other antibiotics) are an attractive option due to their *in vitro* activity (5). However, increasing resistance to SXT has complicated the treatment and resistance determinants such as *sul* and *dfrA* genes, class 1 integrons and mobile genetic elements have been reported to contribute to SXT resistance (6–8). The aim of this study was to establish the link between 4 SXT resistant *S. maltophilia* isolates collected in a neonatalogy unit and to decipher the genetic basis of resistance to SXT.

## MATERIALS AND METHODS

### Study design

The longitudinal study was conducted in the neonatal intensive care unit in CENHOSOA hospital in Antananarivo, Madagascar (08/27/2014–03/06/2015) and was already described in Bonneault *et al*. 2019 (9). Briefly, 22 newborns (NBs) were included in the cohort and were followed until discharge or death. Average unit stays lasted 18 days. All health-care workers (HCWs) and NBs’ accompanying family members (FMs; usually the mother, involved in the basic infant care, except for one child who had four distinct accompanying FMs) were also followed. In total 22 NBs, 21 HCWs and 24 FMs were included in the study. At enrollment, a rectal swab was obtained from the NB and a stool sample from the FM to detect E-ESBL colonization. Rectal swabs were systematically obtained from the NB on a weekly basis. For stays <7 days in the unit, a stool sample was obtained the day of discharge. Stools were also collected from the FMs and from the HCWs every week. Environmental swabbing was performed at the beginning, middle and the end of the investigation. The study was approved by the Madagascar Public Health Ministry Ethics Committee (Reference number: 040– MSANP/CE).

### Bacteriological analyses

All samples were cultivated on CHROMagar™ ESBL (CHROMagar, Paris, France) and each colony morphotype was identified by mass spectrometry (MS) MALDI-TOF (Bruker Daltonics, Bremen, Germany). Antimicrobial susceptibility testing was performed on each isolate according to the standard disc methods described in the 2018 CASFM guidelines. In this study, we only focused on *S. maltophilia* positive samples.

### Whole genome sequencing (WGS) & Bioinformatic analysis

DNA extraction was performed on 5 mL of liquid cultures grown overnight at 37°C in a Luria Bertani infusion medium by using the Cador Pathogen Extraction Kit (Indical Bioscience) on the Qiacube HT (QIAGEN, France) device according to the manufacturer’s protocol for Gram-negative bacteria. DNA quantity and purity was assessed by using Nanodrop 2000/200C (Thermo Fisher Scientific, Waltham, MA, USA). Library preparation was conducted by using the Nextera XT DNA Sample Kit (Illumina, San Diego, CA, USA). WGS was performed on a NextSeq 500 platform (Illumina) by using 2 × 150-bp runs. FqCleaner version 3.0 was used to eliminate adaptor sequences (10, 11), reduce redundant or overrepresented reads (12), correct sequencing errors (13), merge overlapping paired reads, and discard reads with Phred scores (measure of the quality of identification of nucleobases generated by automated DNA sequencing) <20. Illumina reads *de novo* assembly was performed using Spades (14). Acquired resistance genes were detected by the resfinder software (15). The genomes were annotated by using prokka and PATRIC web server (16, 17). The sequence types (ST) were determined *in silico* with the public database for *S. maltophilia* (https://pubmlst.org/smaltophilia/). Phylogenetic analysis based on whole genome sequences was done using the Parsnp program from the harvest suite, gubbins and RaxML (18–20). Mutation detection in SXT resistant *S. maltophilia* strains was performed with the Breseq 0.31.1 software (*S. maltophilia* D457 was used as reference genome) (21). All non-synonymous mutations were analyzed with PROVEAN (Protein Variation Effect Analyzer) (13) to predict the functional deleteriousness caused by the missense mutation. Plasmid detection, typing and reconstruction were performed with the MOB-suite software (22).

### Predicted structure and molecular docking

The secondary and tertiary structures of proteins were predicted by using the Raptor X server (23–26). The wild-type amino acid sequence from the D457 strain was chosen as template for homology modeling. We used autodock vina to calculate Gibbs free energy of binding (ΔG_bind_) of ligands (trimethoprim) with targets. They were further converted to the predicted inhibition constants (Ki_pred_) with this formula (27):

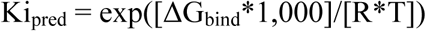

where R (gas constant) is 1.98 cal(mol*K)-1, and T (room temperature) is 298.15 Kelvin.

### Quantification of biofilm formation

The biofilm assay was performed as previously described (28) but with slight modifications. Overnight cultures of *S. maltophilia* in 5 mL Luria Bertani infusion medium reaching an optical density at 620 nm equivalent to 1 (OD620) (approximately 1*10^9^ CFU/mL) were transferred to the wells of a sterile flat-bottomed 96-well polystyrene microtitre plate and incubated for 24h at 35°C +/-2°C. Non adherent cells were subsequently removed by washing twice with 200 µl of sterile distilled water. The amount of biofilm biomass was assessed by crystal violet staining. Biofilms were stained with 125 µl of 1.0 % (w/v) crystal violet for 15 min. The dye solution was discarded, and the plate was washed three times with sterile distilled water and allowed to air-dry for 24 h at room temperature. Stained biofilms were exposed to 30.0 % (v/v) acetic acid for 15 min, and the OD 620 of the extracted dye was subsequently measured. The average OD values was calculated for all tested strains and negative controls, since all tests are performed in triplicate and repeated three times. Second, the cut-off value (ODc) was established. It is defined as three standard deviations (SD) above the mean OD of the negative control: ODc=average OD of negative control + (3×SD of negative control). OD values of a tested strain are expressed as average OD value of the strain reduced by ODc value (OD=average OD of a strain -ODc). ODc value is calculated for each microtiter plate separately. Based upon the previously calculated OD values: OD ≤ODc = no biofilm producer; ODc ≤OD ≤2 × ODc = weak biofilm producer; 2 × ODc≤OD≤4×ODc = moderate biofilm producer; 4ODc<OD = strong biofilm producer. All tests were performed in triplicate and repeated three times.

## RESULTS

### Epidemiological data

During the cohort conducted in the neonatal unit - (see Materials and Methods), 4 *S. maltophilia* were isolated. A first *S. maltophilia* strain (MS MALDI-TOF score of 2.736) was isolated in January, 07th 2015 from an environmental swab (476SM) realized on a water tap out of the three sinks present in the neonatalogy unit. Fourteen days later a second *S. maltophilia* strain (MS MALDI-TOF score of 2.120) was isolated from a rectal swab of a premature newborn (517SM) admitted to the neonatal ICU. Subsequently, two strains of *S. maltophilia* were isolated from the fingerprints of 2 nurses at day 47 (629SM) and 56 (646SM) after the first isolation from the water tap (MS MALDI-TOF scores of respectively 2.102 and 2.052) (Table 1).

**Table 1:** Antimicrobial susceptibility testing of the four *S. maltophilia* strains. Abbreviations: SXT: trimethoprim-sulfamethoxazole, TCC: Ticarcillin-clavulanate, CAZ: ceftazidim, LVX: levofloxacin, MNO: minocycline, MEM: meropenem, ETP: ertapenem and IPM: imipenem.

| Strains | Origin | Isolation date | Sample | inhibition zone diameters (mm) |  |  |  |  |  |  |  |
| --- | --- | --- | --- | --- | --- | --- | --- | --- | --- | --- | --- |
|  |  |  |  | SXT | TCC | CAZ | LVX | MNO | MEM | ETP | IPM |
| 476SM | Watertap | January 07, 2015 | Environmental swab | 6 (R) | 35 (S) | 22 (S) | 21 (I) | 25 (S) | 6 (R) | 6 (R) | 6 (R) |
| 517SM | Baby | January 21, 2015 | Rectal swab | 6 (R) | 37 (S) | 23 (S) | 21 (I) | 27 (S) | 6 (R) | 6 (R) | 6 (R) |
| 646SM | Nursing staff1 | February 23, 2015 | Fingerprint | 6 (R) | 36 (S) | 23 (S) | 21 (I) | 24 (S) | 6 (R) | 6 (R) | 6 (R) |
| 629SM | Nursing staff2 | March 04, 2015 | Fingerprint | 6 (R) | 34 (S) | 21 (S) | 21 (I) | 26 (S) | 6 (R) | 6 (R) | 6 (R) |

### Antimicrobial susceptibility testing

All 4 isolates were resistant to cotrimoxazole (SXT), ertapenem (ETP), meropenem (MEM) and imipenem (IPM), intermediate to levofloxacin (LVX) and sensitive to minocycline (MNO), ceftazidime (CAZ) and ticarcillin-clavulanate (TCC). The antimicrobial susceptibility testing results of the 4 strains are showed in table1.

### Phylogenetic analysis and multilocus sequence-types

A phylogenetic analysis based on whole genome sequences was performed using the Parsnp program. The 4 isolates are identical and clustered with a *S. maltophilia* (LQQS01000001) originating from Malaysia (figure 1). The sequence types (ST) of our four isolates were determined with MLST *in silico* with the public database (https://pubmlst.org/smaltophilia/). They all belonged to a new ST: ST-288.

**Figure 1:**
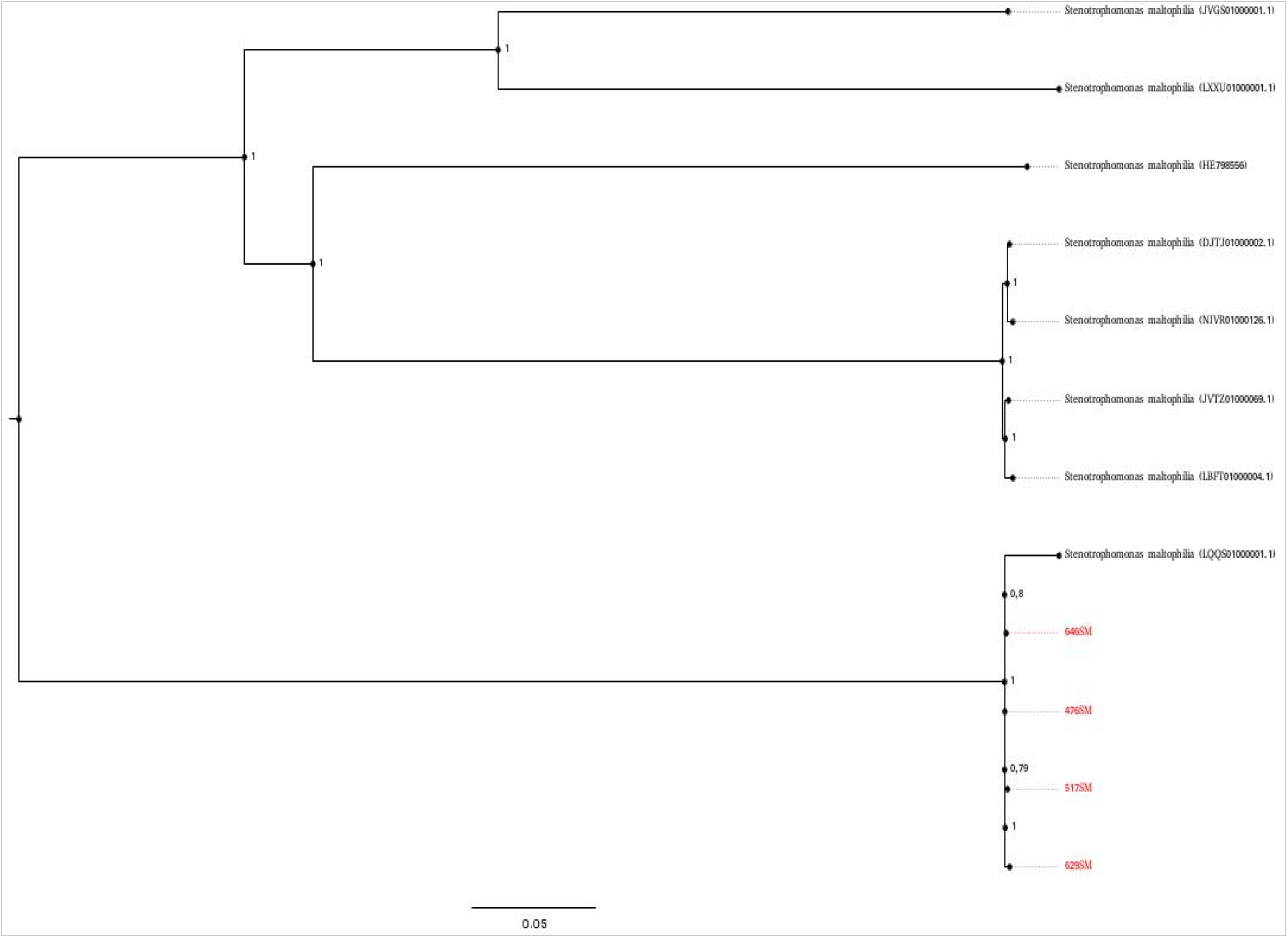
Phylogenetic tree based on coregenome alignment of four *S. maltophilia* including representative genomes from NCBI genbank. Sequences were aligned using ParSNP, genetic recombinaison were removed using Gubbins and phylogenetic inferences were obtained using the maximum likelihood method within raxML software. Bootstrap values are expressed by decimal of 1 000 replicates with a parameter test and shown at the branching points. The branches of the tree are indicated by the genus and species name of the type strains followed by the NCBI gene accession numbers. The four *S. maltophilia* isolates from this study are represented in red color.

### Resistome analysis

Multiple antimicrobial, heavy-metal and arsenic resistance markers were identified in the chromosome of each isolate (Table 2). The resistome of these 4 strains revealed the presence of 27 antibacterial-resistant genes using the resfinder software and PATRIC webservers. The isolates possess two different types of β-lactamases, i.e., Amber class A genes (L2 family), Amber class B metallo-β-lactamase (MBL) genes (L1 family), and five aminoglycoside inactivation enzymes: *aph(2’’)-ia, aph(3’’)-i, aph(3’)-ii/aph(3’)-xv, aph(6)-ic/aph(6)-id*. We also identified 9 efflux pumps conferring antibiotic resistance and two regulators modulating expression of antibiotic resistance genes

**Table 2:**
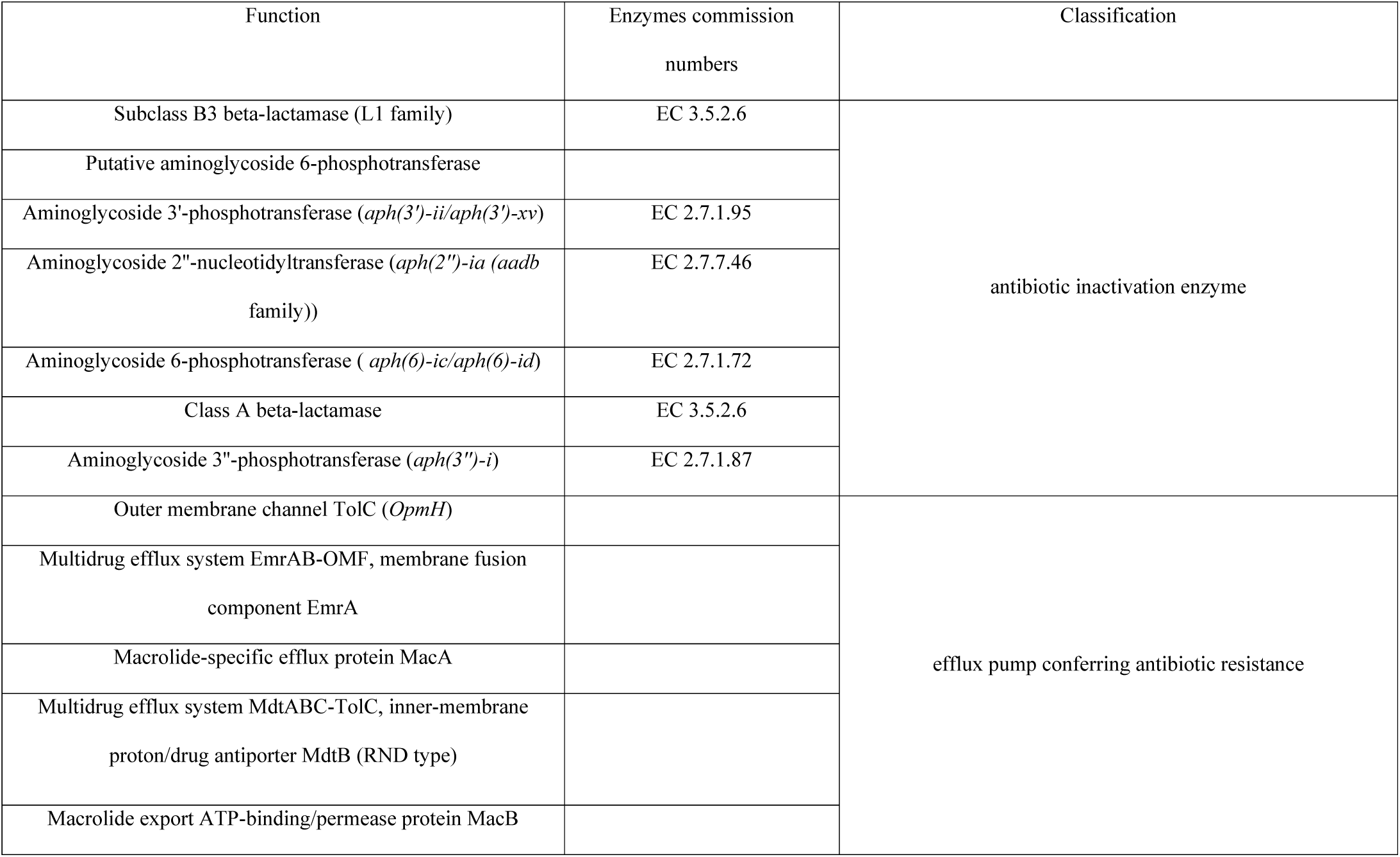
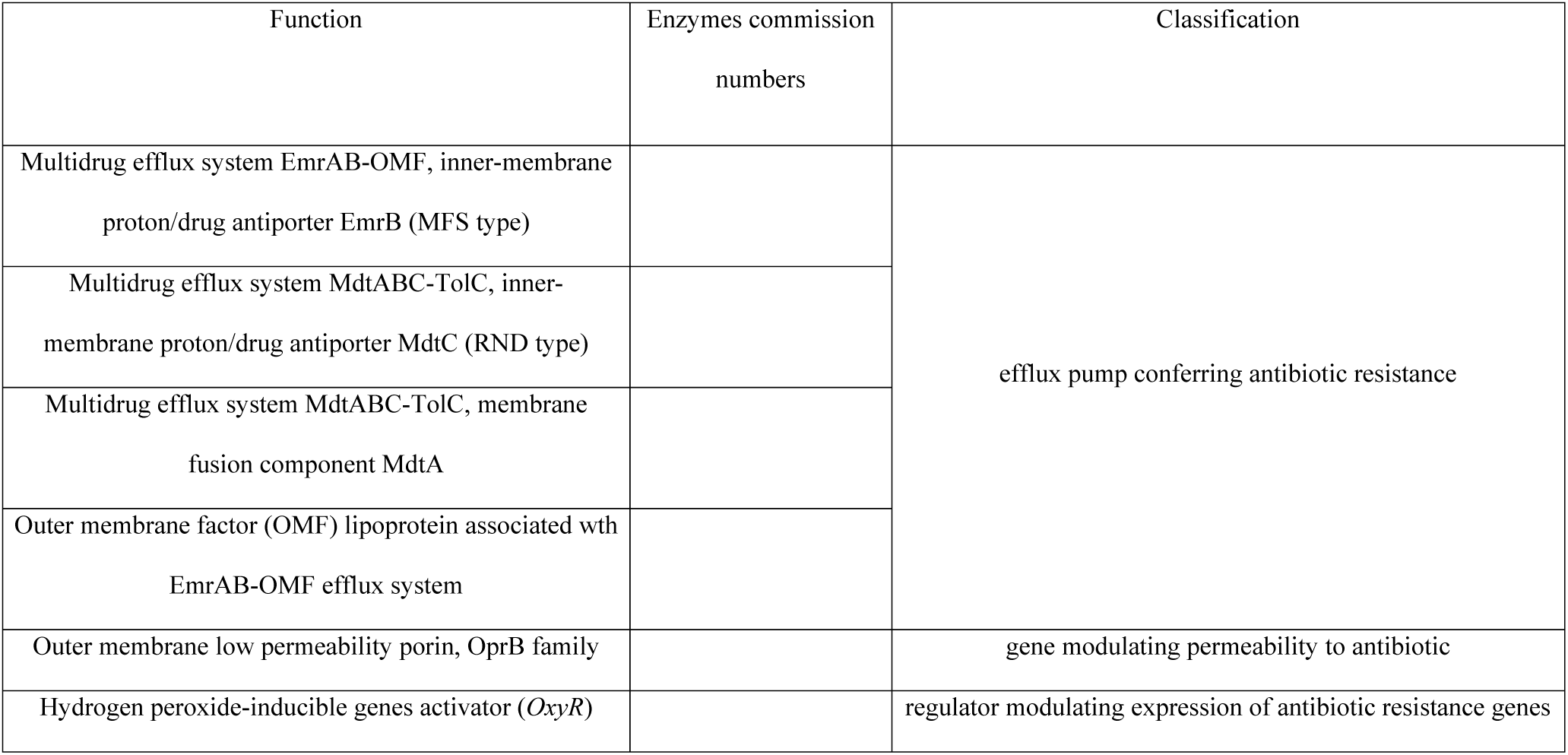
Resistome of *S. maltophilia* isolated in the neonatology unit.

### Plasmid analysis

Mobsuite software enabled us to identify and reconstruct two plasmids: an IncP conjugative plasmid harboring the aminoglycoside 6-phosphotransferase gene (*sph* gene) and which presented 86% nucleotidic identities with the plasmid p35734 found in *E. cloacae*. Moreover we founda non-transferable plasmid with a RND efflux transporter conferring resistances to cobalt, zinc, cadmium and which presented 99% nucleotidic identities with the plasmid pLMG930 found in *Xanthomonas euvesicatoria*. As we used the short-reads sequencing technique, we were unable to circularize the plasmids.

### Genetic environment of resistance genes

All isolates carried class I integrons that contains *sul*1, *aad*B, *aph*3, *aph*6 genes and the entire heat shock protein gene *groEL* upstream the *sul1* gene (Figure 2).

**Figure 2:**
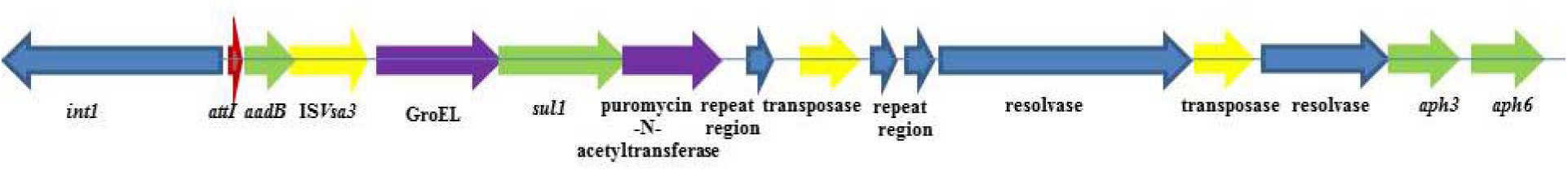
Structure of class I integrons in the *S. maltophilia* isolated in the neonatology unit.

### Mutations and resistance to SXT

When using *S. maltophilia* D457 as reference genome a total of 327 deletions, 286 insertions and 2133 substitutions were detected with the Breseq 0.31.1 software. In all isolates, single nucleotide mutations in genes encoding enzymes involved in folic acid metabolism which can be inactivated by trimethoprim were detected in *folA* (A114T:GCC→ACC), *folC* (L339V:TTG→GTG) and *folM* (A72P:GCC→CCC) which encode respectively the dihydrofolate reductase, dihydrofolate synthase and FolM alternative dihydrofolate reductase 1. Only the mutation in *folA* has an effect on the activity of the protein as assessed by PROVEAN analysis

### Predicted structure and molecular docking

The Gibbs free energy of binding (ΔG_bind_) between the FolA protein (mutated and wild-type) and trimethoprim was calculated and values of - 6.9 kcal/mol and −7.1 kcal/mol were found for the mutated and the wild-type proteins, respectively. The predicted inhibition constants (Ki_pred_) between trimethoprim and the FolA wild-type protein was 3.59×10^-6^ and 8.39X10^-6^ between trimethoprim and the mutated FolA indicating a reduced susceptibility to trimethoprim.

### Quantification of biofilm formation

The amount of biofilm biomass was assessed by crystal violet staining as explained in table 4. All 4 isolates had an average OD_650nm_ value of 0.145 (ODc= 0.086), meaning they are weak biofilm producers. Additionally, we detected in the four isolates modifications in genes involved in biofilm production: 43 substitutions in the rmlA gene, 32 substitutions in the spgM gene, two insertions and three substitutions in the intergenic region between the manA and spgM genes, two substitutions in the intergenic region between the rfbB and rmlA genes and one substitutions in the intergenic region between the rpfC and rpfF genes. These genetic modifications are probably the cause of weak biofilm production. The list of all mutations detected is showed in supplementary material S1.

**Table 4:** Optical density (650nm) of biofilm formation of four strains.

| ID strain | 517SM | 476SM | 629SM | 646SM | Standard deviation | OD<sub>negative</sub> |
| --- | --- | --- | --- | --- | --- | --- |
| test 1 | 0,212125 | 0,11725 | 0,174125 | 0,1245 | 0,00698851 | 0,08659053 |
| test2 | 0,174125 | 0,19625 | 0,125375 | 0,100625 | 0,01431034 | 0,08368101 |
| test3 | 0,13575 | 0,135625 | 0,124 | 0,12375 | 0,00484031 | 0,08952092 |
| Average | 0,174 | 0,14970833 | 0,14116667 | 0,11629167 | 0,00871305 | 0,08659749 |

## Discussion

The prevalence of *S. maltophilia* has increased in hospitals worldwide simultaneously with the emergence of a myriad of other antibiotic resistant bacteria (29). Here, we demonstrated, based on WGS phylogeny, the clonal transmission of *S. matophilia* resistant to SXT in a neonatalogy unit, which might be explained by its ecology and fitness in hospitals wards and/or by poor hygiene management. The determination of a new ST is in accordance with the high plasticity and capacity of this bacterium to adapt to specific niches and develop new characteristics. We showed in our study that the same strain of *S. maltophilia* was isolated four times on a period of two months in a neonatal ward of a hospital in Antananarivo, confirming its ability to persist and spread in the medical environment. Selective pressure imposed by specific conditions in a hospital environment could promote the survival of certain STs with an adaptive advantage for this specific setting and lead afterwards to their clonal spread.

The reduced susceptibility of *S. maltophilia* to most antibiotics can be attributed to both intrinsic and acquired resistances. The proteins mediating intrinsic resistance in *S. maltophilia* include chromosomally encoded multidrug efflux pumps such as SmeABC, SmeDEF, SmeYZ, SmeOP-TolCSm, antibiotic-inactivating enzymes (L1/L2 β-lactamases and aminoglycoside inactivating enzymes), and the chromosomally encoded Qnr pentapeptide repeat proteins (30) which are present in most if not all strains of *S. maltophilia*, as in our Madagascan strains, suggesting that they did not arise during the recent evolution of resistance caused by antibiotic therapy. In addition, *S. maltophilia* can acquire mechanisms to increase its resistance pattern through horizontal gene transfer via plasmids, and subsequently by recombination processes with integrons, transposons and genomic islands (GIs).

The four isolates, which are genotypically identical, were resistant to SXT, one of the therapeutic choices. The resistance of Gram-negative bacteria to sulfonamides is mainly conferred by the acquisition of either *sul1* or *sul2*, encoding dihydropteroate synthases (31). The *sul1* gene carried by class 1 integrons and sul2, which is linked to insertion sequence common region (ISCR) elements, was identified in SXT-resistant *S. maltophilia* isolates (5, 32, 33). The resistance to trimethoprim in *S. maltophilia* is mainly conferred by the dihydrofolate reductase *dfr* genes such as *dfrA1, dfrA5, dfrA12, dfrA17*, and *dfrA27* which are usually located within class 1 integrons as part of various resistance gene cassettes. Both types of *sul* and *dfr* genes can occur together in high-level SXT-resistant isolates (6, 34). Moreover, the efflux pumps SmeDEF, TolCsm, and SmeYZ are associated with SXT resistance (35–37). We found that the *sul1* gene was present but no *dfr* genes were detected, pointing out the presence of another mechanism of resistance to trimethoprim. We were able to show that a point mutation (A114T:GCC→ACC) in the dihydrofolate reductase gene (*folA*) decreased the affinity of trimethoprim to the FolA protein ensuring therefore resistance to this antibiotic.

We have also shown the presence of two plasmids, one of which harbored heavy metal resistance genes and which was almost identical to the pLMG930 plasmid found in *Xanthomonas euvesicatoria*, a bacterial spot-causing xanthomonads. This indicates a high probability of dissemination of the strain in different ecological niches, notably those contaminated with heavy metals. It is well known that the presence of both metals and antibiotic resistance genes play a major role in the persistence, selection and spread of antibiotic- and metal-resistant bacteria in anthropogenic environments heavily contaminated with detergents, heavy metals and other antimicrobials [29, 30]. In developing countries, rivers, lakes and lagoons are often contaminated with untreated hospital and industrial effluents and also by urban storm-water containing anthropogenic pollutants due to intensive uncontrolled urbanization. These are optimal conditions for bacterial development and the spread of antibiotic-resistant bacteria.

Biofilm formation in bacteria is a multifactorial event that depends on surface characteristics, motility of strains, genes involved in biofilm formation, and other factors, and is usually correlated with a higher level of resistance to antibiotics and disinfectants (38). Different factors influence the physiology of biofilm formation in *S. maltophilia*, namely the SmeYZ efflux pump that confers resistance to antimicrobials (37), the iron level in the media (39), and histidine kinase and BfmAK system (40). Interestingly, our study revealed a negative correlation between the simultaneous presence of genes involved in biofilm formation such as *spgM, rmlA* and *rpfF* genes and the biofilm production. This correlation could be due to the mutations identified in the three previous genes. However, a clone was detected along a period of two months in the neonatlogy ward pointing out a persistence of the strain in this environment.

In conclusion, this work represents the first characterization at the genomic level of SXT resistant *S. maltophilia* strains circulating in a neonatology ward of a hospital in Antananarivo, Madagascar. We also pointed out the possible role of a point mutation in the *folA* gene conferring resistance to trimethoprim. Clonal relatedness between strains indicated the transmission and the persistence of *S. maltophlia* in the hospital setting and the threat it could represent for newborns, especially for preterms.

## Data availability

These whole genome shotgun projects have been deposited at DDBJ/ENA/GenBank. under the accession <u>VFJF00000000</u>, <u>VFJG00000000</u>, <u>VFJH00000000</u> and <u>VFEX00000000</u> corresponding respectively for the strains.517SM, 629SM, 646SM and 476SM.

## Acknowledgments

We would like to thank Tatianah Seheno Rivomanantsoa, the field investigator, and the staff of pediatric and neonatalogy units’ at the CENHOSOA Hospital, Antananarivo, Madagascar. We would also like to thank the program “Actions concertées inter-pasteuriennes” (ACIP: grant no. A-22-2013) which supported this work and staff of the “Plateforme de Microbiologie Mutualisée (P2M)” at Institut Pasteur Paris where the whole genome sequencing was performed.

